# Coupling Wright-Fisher and coalescent dynamics for realistic simulation of population-scale datasets

**DOI:** 10.1101/674440

**Authors:** Dominic Nelson, Jerome Kelleher, Aaron P. Ragsdale, Gil McVean, Simon Gravel

## Abstract

Coalescent simulations are widely used to examine the effects of evolution and demographic history on the genetic makeup of populations. Thanks to recent progress in algorithms and data structures, simulators such as the widely-used `msprime` [1] now provide genome-wide simulations for millions of individuals. However, this software relies on classic coalescent theory and the corresponding assumptions that sample sizes are small relative to effective population size and that the region being simulated is short. Here we show that coalescent simulations of long regions of the genome exhibit large biases in identity-by-descent (IBD), long-range linkage disequilibrium (LD), and ancestry patterns, particularly when sample size is large. We present a Wright-Fisher extension to `msprime`, and show that it produces more realistic distributions of IBD, LD, and ancestry proportions, while also addressing more subtle biases of the coalescent. Further, these extensions are more computationally efficient than state-of-the-art coalescent simulations when simulating long regions, including whole-genome data. For shorter regions, efficiency and accuracy can be maintained via a flexible hybrid model which simulates the recent past under the Wright-Fisher model and uses coalescent simulations in the distant past.

## 2 Introduction

Coalescent simulations are widely used in the development of statistical tools for genetics research [2–7]. They generate large sequence datasets which can be used to test our understanding of evolution or the ability of statistical tests to identify disease variants and avoid confounding factors. Coalescent theory has been used extensively for this purpose, with Hudson’s ms simulation program [8] having been cited over two thousand times since its publication in 2002.

The more recent `msprime` coalescent simulation software [1] implements Hudson’s original algorithm [9], but with a performance increase of several orders of magnitude. This is achieved largely through the introduction of a new data structure, the succinct tree sequence [10, 11], which is extremely efficient at storing genetic variation, and the genetically relevant component of the genealogy of large numbers of samples. For example, simulating a 100 megabase region in a sample of 100,000 individuals generates an 88MB uncompressed succinct tree sequence, whereas the Newick tree format used by `ms` takes approximately 3.5TB of space [1].

Simulations are useful to the extent that they accurately reflect genetic variation within a real sample having undergone a similar demographic history. However, the coalescent is known to be biased relative to the Wright-Fisher model when the sample size is large [12] or for events in the recent past [13]. However, these biases have had limited practical impact because collecting such large empirical data sets was prohibitively costly and the simulation of such large samples was computationally overwhelming. Both limitations have now been lifted: sequencing datasets now regularly include thousands of sequenced genomes, and `msprime` can simulate hundreds of thousands of genomes on a laptop computer. The assumptions of the underlying coalescent models should be carefully reexamined in this context.

We highlight qualitative and quantitative inaccuracies in coalescent simulations of long regions, due to violated assumptions of the underlying genealogical model. We implement an extension to `msprime` which corrects the majority of these biases while reducing computational cost relative to coalescent simulations for large samples and long regions. This is accomplished by implementing a backwards-in-time Wright-Fisher model within `msprime`, which generates biologically plausible genealogies regardless of sample size. Models combining coalescent and Wright-Fisher dynamics can benefit from strengths of both approaches [13], and our implementation also allows for hybrid simulations which combine the advantages of the Wright-Fisher model in recent generations with the efficiency of the coalescent in the distant past, maintaining high performance for shorter regions when the Wright-Fisher model alone is less efficient.

### 2.1 Motivation

This work was motivated by our observation that large-scale coalescent simulations resulted in unrealistic relatedness among samples, where nearly every pair of samples were second-or third-degree cousins according to the time to their most recent common ancestor. We traced this phenomenon to samples having more than 2^*t*^ simulated ancestors at generation *t* in the past, allowing more recent relatedness than is possible under diploid inheritance. This is a side-effect of Hudson’s coalescent algorithm lacking a diploid population pedigree [12].

Hudson’s coalescent model assumes a small region being simulated [14], and so does not account for multiple simultaneous recombinations during meiosis. Since the time between events in Hudson’s model can be less than a single generation, the per-generation recombination rate in long genomic regions is maintained by multiple single recombinations, but these occur through independent events that involve more than two parental lineages. This is unrealistic, and biases diversity predictions. An independent lineage is created with each event, leading to a large number of new lineages in the timescale of the first generation (Figure 1b).

**Figure 1:**
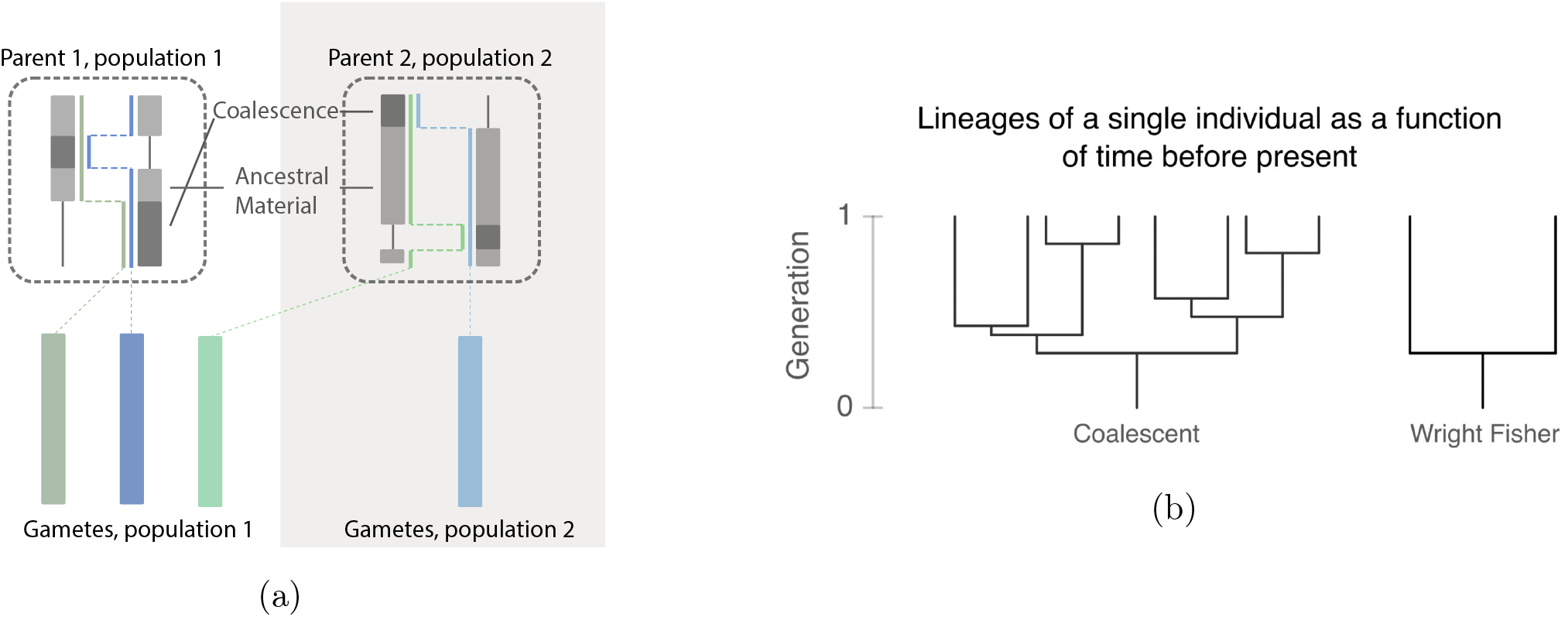
Comparing coalescent and Wright-Fisher lineages. (a) A single backwards-in-time Wright-Fisher generation in our implementation. First, migration moves sample lineages (bottom) between populations. Lineages then draw parents (dotted outlines), and are split by recombination events into the two parental copies of the genome. Each parental copy is then merged into a new lineage, with coalescence events occurring where sample regions overlap (shaded areas). These new lineages are carried on to the next generation. (b) Schematic of lineages simulated under different models over a region of length 8 Morgans.

This peculiarity of Hudson’s coalescent is negligible in small samples without migration, because simultaneous recombinations have a small effect on the simulated genealogies [13, 15]. In larger samples, or under migration models, recent events induce long-range correlations along the genome due to the diploid population pedigree, and these are not accounted for by Hudson’s coalescent [12].

At whole-genome scale this becomes particularly clear. To highlight the magnitude of the genealogical distortions which can occur, we first use both the coalescent and Wright-Fisher models to simulate 10,000 haploid samples in a diploid population of constant size 10,000. Each sample contains 22 chromosomes of realistic lengths. Figure 2 shows that the number of lineages in the coalescent simulation increases very rapidly to reach 10 times the haploid population size 2*N* (A similar result was shown in [16]). In the Wright-Fisher simulation, the initial growth in number of lineages is much slower and can never exceed those present in the population. We describe some ways these differences affect variation within simulated cohorts in the Results section.

**Figure 2:**
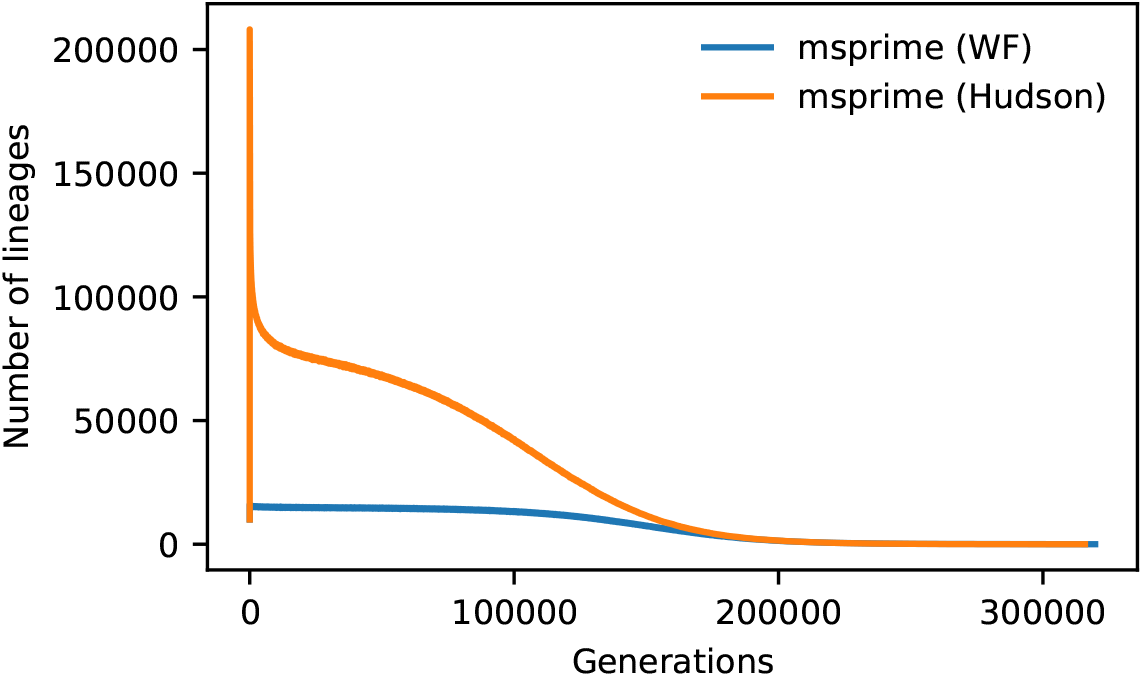
Number of surviving lineages over time in coalescent and back-in-time Wright-Fisher dynamics. We simulated 10,000 haploid whole genomes with 22 chromosomes of realistic lengths in a population of 10,000 diploid individuals. The method for simulating multiple chromosomes is described in Section 4.1.

While genealogical distortions are most clear in the first few generations, this explosion of lineages also affects genealogies in the more distant past. Figure 2 also shows that, despite rapid coalescence lowering the initial spike in the number of lineages, their number remains above the population size until 150,000 generations in the past.

## 3 Materials and Methods

### 3.1 Implementation

Implementing a discrete backwards-in-time Wright-Fisher model within `msprime` is conceptually straightforward. To understand the modifications needed, we first outline Hudson’s original algorithm to simulate samples under the coalescent model. This brief overview is intended to give context to the modifications which enable Wright-Fisher simulations to be performed in the same framework. More details of how Hudson’s algorithm is implemented in `msprime` are given in [1].

First, a number of randomly-mating populations are specified, including effective sizes and migration rates over time. Samples are introduced as haploid lineages within the populations, and the region of the genome being simulated is specified. The algorithm then constructs the genealogy of each locus within this region by tracing its lineages backwards in time and tracking genomic segments that are ancestral to the sample.

To begin, each lineage contains a single ancestral segment spanning the whole simulated genomic region of a sample. As time proceeds backwards, lineages can be split by recombination events (leaving the amount of ancestral material unchanged), or participate in common ancestor events, where any overlapping regions coalesce (reducing the amount of ancestral material). The recombination rate depends on the sum of the genetic map distance spanned by ancestral segments carried by all extant lineages, and common ancestor events occur at a rate determined by the number of uncoalesced lineages and the effective population size. Migration events move haploid lineages between randomly-mating populations, and demographic events modify the number of populations or their size and growth rate parameters. Recombination and common ancestor events are generated at rates depending on the amount of extant ancestral material, and the simulation terminates when every position on the genome has a most recent common ancestor

Implementing a back-in-time Wright-Fisher model requires two important changes to Hudson’s algorithm. First, rather than drawing a time to the next event from an exponential distribution, we iterate though discrete generations and draw the events which occur at each time. Second, we modify the way recombination events are carried out, to account for the possibility of multiple recombinations in a single transmission: we model the number and spatial distribution of breakpoints as a Poisson process, with rate equal to the per-generation recombination rate (i.e., the distance in Morgans). Crossover interference and non-crossover tracts would be straightforward to implement but are not included here. An overview of this model is illustrated in Figure 1a and the detailed order of events occurring at each generation is given in the Supplement. This model ensures that each gamete has a unique diploid parent.

## 4 Results

In this section, we first highlight qualitative differences between the coalescent and backwards Wright-Fisher models, and we show that the Wright-Fisher models provide a better description of the data while increasing tractability. We also study the impact of our implementation on previously-studied differences between coalescent and Wright-Fisher models, such as differing proportions of singletons and doubletons [13]. These are shown in Figure S1 and Table S1. We focus here on large discrepancies which, to our knowledge, have not been previously examined.

### 4.1 Distribution of IBD

Under the Wright-Fisher model, diploid inheritance constrains the possible gene genealogies [12] and introduces correlations in IBD sharing along long simulated regions: two samples with a recent common ancestor may be IBD at several distant regions of their genome (for example on different chromosomes). In the coalescent, gene genealogies of unlinked loci are constructed independently [12]. This is a poor model for close relatives, since it is unlikely that any two samples will share more than one IBD segment from the same common ancestor, however recent.

Modelling relatedness patterns is important in large cohorts, where cryptic relatives are common [17, 18]. To illustrate the significance of explicitly modelling diploid inheritance in a sample with close relatives, we simulated 500 human haploid whole genomes (chromosome lengths and recombination rates are described in the Supplement) in a diploid population of constant size 500 under the coalescent and Wright-Fisher models. We simulate a modestly-sized cohort since the effects we wish to highlight depend only on the ratio of sample size to effective population size, and since it simplifies analysis of the resulting data. Larger cohorts can be simulated very efficiently, as shown in Section 4.3.

We used the simulated genealogies to extract IBD segments inherited from common ancestors no more than 10 generations in the past. Closer relatedness means more IBD segments and longer average length, leading to a relationship between number of segments and total length of IBD which is typically used in identifying relative status [17]. Figure 3 shows the qualitative difference between the two models, with the coalescent model exhibiting far too few IBD segments for closely related samples and poor clustering by TMRCA. By contrast, the Wright-Fisher model shows the expected qualitative distribution of relatedness [17]. An analytical model for the expected number and length of shared ancestry segments (shown as black dots in Figure 3) is provided in the Supplement.

**Figure 3:**
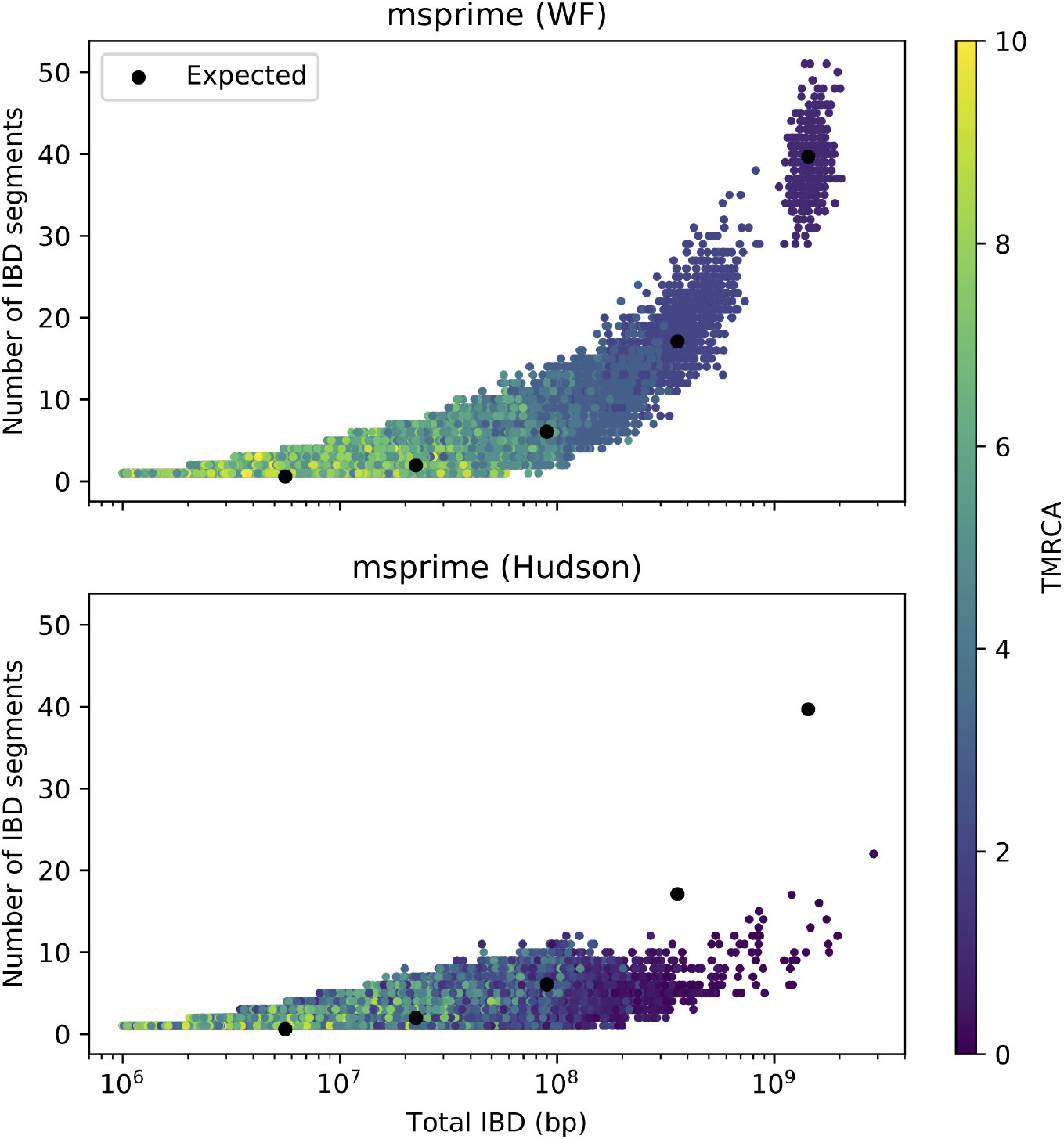
Number of IBD segments between pairs of individuals versus total length of shared IBD segments. 22 chromosomes of realistic lengths, simulated under Wright-Fisher model (top) and coalescent (bottom), compared to expectation under Equations (1) and (2). 500 haploid samples simulated with a diploid population size of 500.

The long-range correlations induced by genealogical relatedness can also be measured as linkage disequilibrium between distant loci. This LD is used to estimate sizes of small populations in conservation genetics [19]. Hudson’s coalescent does not capture such LD patterns, whereas the Wright-Fisher extension to `msprime` predicts the patterns of LD expected under diploid mating (Figure S2).

### 4.2 Ancestry Variance Following Admixture

In admixed populations, simulations should capture patterns of ancestry variation among present-day samples. The distribution of ancestry within recently admixed populations can be strongly dependent on pedigree structure, making coalescent simulations of these scenarios problematic.

We consider the variance of ancestry proportions following a single pulse of migration. Ancestry variance can be divided into genealogical variance and recombination variance [20]. In the first few generations after admixture, variance is driven by genealogical differences in the number of migrant ancestors of each individual. As time goes on, each present-day individual has more ancestors from the admixed generation, exponentially reducing this source of variance. After roughly 10 generations, variation in the amount of genetic material received from each migrant ancestor becomes a stronger source of variance [20].

We performed whole-genome simulations to evaluate how well the Wright-Fisher and coalescent models capture variance in ancestry. Figure 4 shows ancestry variance from simulations of 80 haploid samples in a diploid population of size 80, and a single event of 30% admixture at varying time in the past. These parameters were chosen to match those in [20], and the qualitative results depend only on the ratio of sample size to effective population size. The approximate expected values are derived from an argument similar to the one presented in the supplement for IBD sharing and outlined in [20].

**Figure 4:**
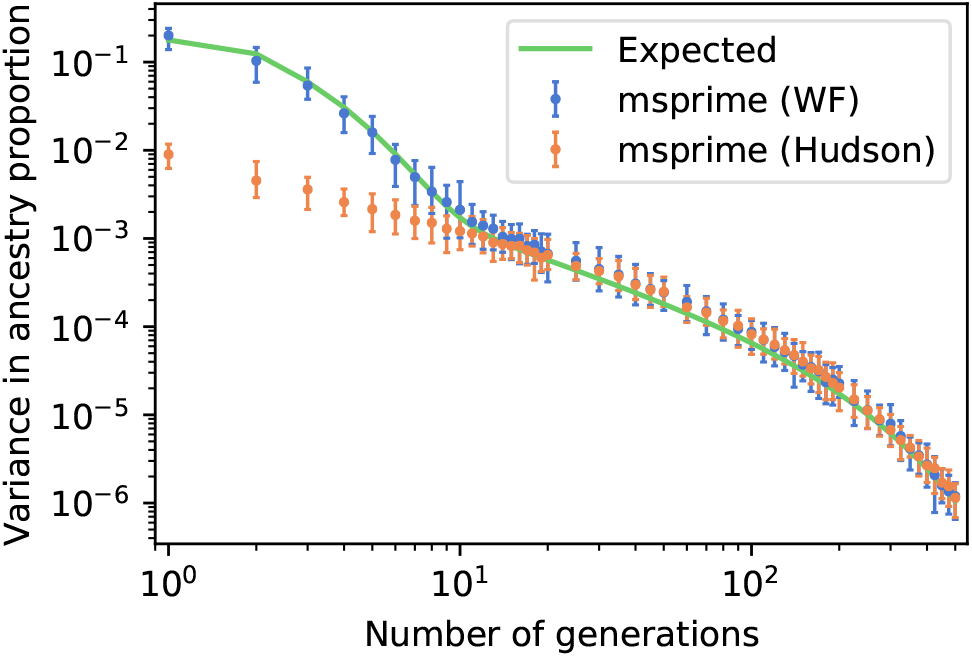
Variance in ancestry after a single admixture event, as a function of time since admixture. Calculated from 80 haploid samples in a diploid population of size 80, with 30% admixture proportions. Error bars show 95% confidence intervals over 50 simulations.

The Wright-Fisher model captures both short- and long-term variance in ancestry, as expected. In the coalescent simulations the initial phase of genealogical variance is not present, leading to a 20-fold underestimate of the variance in ancestry. Lacking a diploid population pedigree, whole-genome coalescent simulations of recently admixed populations do not reflect the distribution of ancestry expected in a large cohort.

### 4.3 Performance

The main advantage of `msprime` over alternate simulators is speed and scalability. This is achieved by efficient algorithms and, especially, new data structures for storing and manipulating ancestral states throughout a simulation. We therefore need to ensure that the present modification preserves these advantages.

An important part of `msprime`’s efficiency is that Hudson’s algorithm simulates only those events which affect variation within the samples. This means that long stretches of time can be traversed in a single step if they contain no such events. However, Figure 2 shows that the number of lineages in whole-genome coalescent simulations is so high that the time between events will on average be much less than a single generation. Furthermore, these lineages come at an additional memory and computational cost for the coalescent model.

Our Wright-Fisher extension is integrated with `msprime`’s core simulation framework, and can also easily be combined with coalescent simulations as part of a hybrid model (a hybrid Wright-Fisher/coalescent approach has also been proposed in [13]). This allows us to combine the strengths and efficiencies of both models, using the Wright Fisher for robust recent genealogies, and switching to the coalescent in the deeper past when the number of events per generation is low.

Figure 5 shows a significant performance advantage for pure Wright-Fisher simulations at whole-genome scale, and a modest advantage for slightly shorter regions. For simulations of a small number of chromosomes, the hybrid and coalescent models have nearly identical performance, but the hybrid simulations yield more realistic recent genealogies. Wright-Fisher simulations are therefore faster and provide more biologically plausible outcomes than the pure coalescent approach.

**Figure 5:**
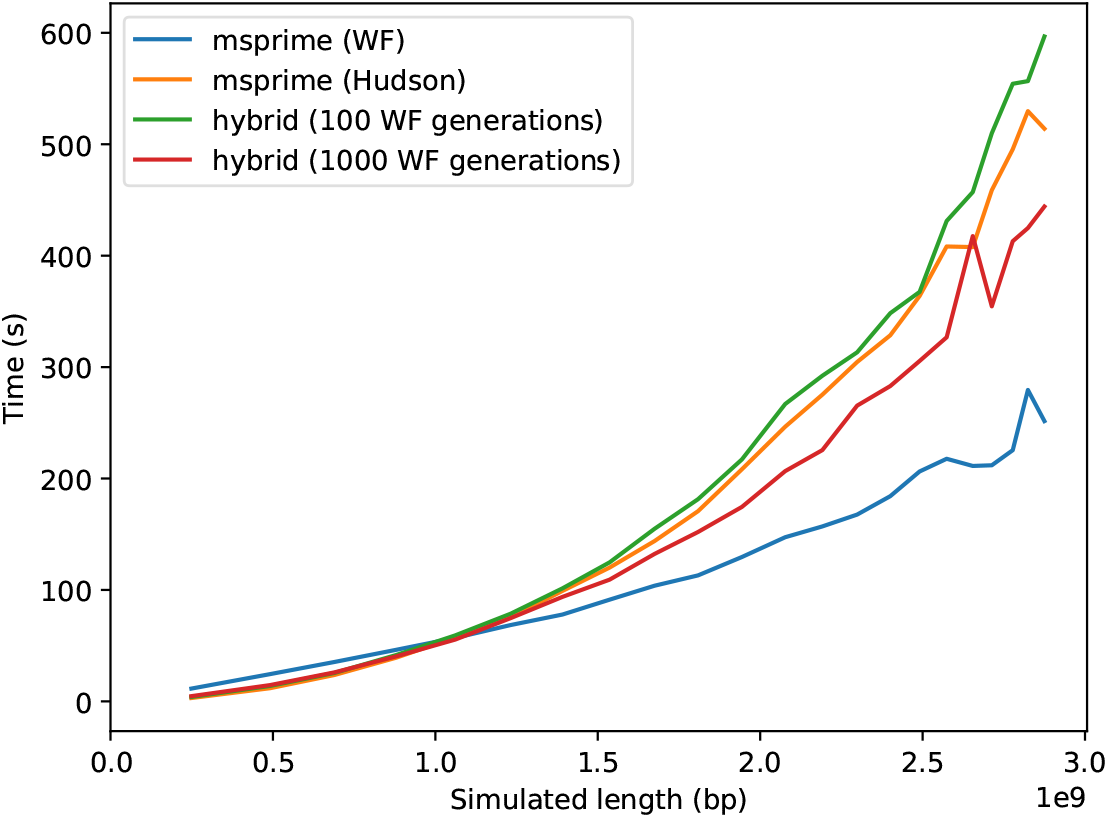
Computation time of Hudson coalescent, Wright-Fisher, and hybrid models. Hybrid models used 100 and 1000 Wright-Fisher generations before switching to the coalescent. Simulations contain from 1 to 22 chromosomes of realistic lengths (using the method described in Section 4.1) in 1000 haploid samples within a diploid population of size 10000. Results for other population sizes are shown in Figure S3.

## 5 Discussion

While the Wright-Fisher model may generate a more realistic pedigree than the coalescent model in the recent past, it is still a highly idealized model. Our implementation does not track monogamous couples, for example, and therefore will overestimate the prevalence of half-sibs and underestimate full sibs compared to a realistic human cohort. Assortative mating and inbreeding are not accounted for, and the migration model, while biologically plausible, is a simplification of the real migration process (see implementation details in Supplement). Care should be taken in applications which are particularly sensitive to fine-scale mating or migration patterns.

Many of these issues can be addressed by allowing simulations to take place within a pre-specified pedigree, which is a natural extension to our backwards-in-time Wright-Fisher implementation. Rather than drawing genealogical links at random according to demographic parameters, lineages can simply follow a known pedigree. When reaching a pedigree founder, simulations can then continue by reverting to either the Wright-Fisher or the coalescent models. Real pedigrees of any size could then be used, from extended families up to population-scale [21], or they could be generated with the desired patterns of monogamy or assortative mating in a separate step. We leave such an implementation for future work.

Other improvements are also possible. Assigning sexes to parents would allow simulation of the X-chromosome and sex-biased migration. Recombination can be extended to model crossover interference and sex-biased recombination, which have effects on the distribution of IBD [22], as well as non-crossover events. The performance of the hybrid model could also be improved. If the number of Wright-Fisher generations were chosen optimally, it is likely to be more efficient than pure Wright-Fisher simulations in nearly all scenarios. Better guidelines for finding this optimal value could be developed, or possibly built into the simulation framework itself.

The limitations of the coalescent model have been well-studied, but are generally shown to have modest effects. This is not always the case - we have shown significant qualitative and quantitative biases in whole-genome simulations of large, complex cohorts. Analysis of such cohorts is challenging, and simulations are a valuable tool for evaluating disease associations and the effects of demography in this context. We have presented here an extension to `msprime` which corrects major biases and increases performance at whole-genome scale, allowing simulations to continue supporting modern large-scale sequencing efforts.

## 6 Web Resources

- `msprime` repository: https://github.com/tskit-dev/msprime
- Documentation (including Wright-Fisher simulations): https://msprime.readthedocs.io

## 1 Supplemental Material

### 1.1 Wright-Fisher Implementation Details

We describe here the precise order of events happening at each generation in our implementation of the Wright-Fisher model. In the first (‘current’) generation, samples are initialized as haploid copies of the region to be simulated (which can later be paired to form diploid individuals). Lineages of each sample are then constructed backwards in time, with each generation explicitly simulated as follows:

1. Migration events are drawn according to the forwards-time rates provided, and migrant lineages are moved to their new population. This is equivalent to migration of gametes, as opposed to migration of diploid individuals. A forwards-time event from population *i* to *j* moves a lineage from population *j* to *i* backwards in time.
2. Demographic events are carried out, such as changes to population sizes or growth rates, mass migrations, or bottlenecks.
3. Each haploid lineage draws a diploid parent within its current population.
4. Recombination occurs, with each breakpoint alternately assigning segments to be inherited from one of the two parental copies of the genome (back-and-forth recombination, see Figure 1b in the main text).
5. Segments inheriting from the same parental copy of the genome are merged into a single lineage, with coalescent events recorded in overlapping regions.

When there is a single ancestral lineage at every position in the simulated genome, the simulation terminates.

Our whole-genome simulations are performed with a single chromosome of length 28.75 Morgans and 22 ‘effective’ chromosomes of realistic lengths separated by 0.5 Morgans. This is not exactly equivalent to simulating fully independent chromosomes. However, this should not have a qualitative impact on the analyses considered here.

### 1.2 Long range linkage disequilibrium

For pairs of loci at low recombination distances (*r ≪* 1), it is unlikely for more than a single recombination event to occur in a given meiosis. In this case, the coalescent accurately models LD between linked loci.

For larger recombination distances, the probability of multiple recombinations must be considered, and loci only become unlinked under odd number of recombinations. This has probability 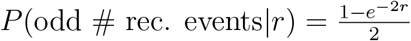, which has a maximum value of 1/2. This leads to non-zero long-range LD, even in the case of fully unlinked loci. The diploid Wright-Fisher captures this, but coalescent estimates decay to zero for increasing *r* (Figure S2).

In diploid individuals, multiple gametes are generated by recombination of the same two parental copies of the genome, and this has consequences on IBD and LD patterns: some classical models of population genetics neglect this and draw pairs of parental chromosomes independently for each gamete ([23]). This also underestimate the amount of linkage disequilibrium among distant sites (Figure S2). Our Wright-Fisher extension to `msprime` therefore correctly accounts for diploid mating (Figure 1b).

### 1.3 An approximate model for IBD sharing

To provide a simple analytical model for this relationship, we examined a pair of haploid samples sharing a single diploid common ancestor at time *t* generations in the past. We derived approximations for the number and length of shared haplotypes given *t*. We can think of the ancestry of each haploid genome as a mosaic formed by copying genomic segments from its 2^*t−*1^ possible ancestors. Similarly, a pair of haploid samples can be seen as a mosaic formed by copying from one ancestor for each sample. We can define paired-ancestry segments as continuous segments having no changes in ancestry in either sample. By this definition, if each sample has *K* chromosomes of total length *L* Morgans, the pair will have on average *K* + 2*Lt* paired-ancestry segments.

Since each haploid sample has 2^*t−*1^ possible ancestors from which to inherit genetic material, a pair of samples will both inherit a paired-ancestry segment from their common ancestor with probability 2^2*t−*2^. Since the ancestor is diploid, they inherit from the same ancestral copy of the genome with probability 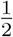. The probability that a paired-ancestry segment is IBD in the pair is therefore 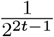, and the expected number of IBD segments *s* between the pair is:

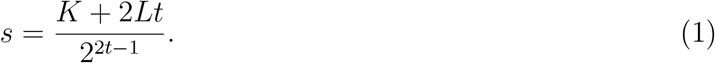

The length of the genome shared, denoted by *x*, corresponds to L times the probability of having a shared ancestor at any particular locus, which is 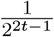, giving:

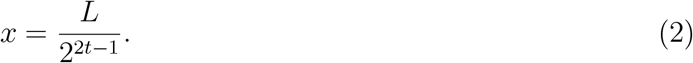

The expected values (*s, x*) are shown in Figure 3 as black dots for *t* from 1 to 5 generations, corresponding to half-siblings, first cousins, and so on.

**Figure S1:**
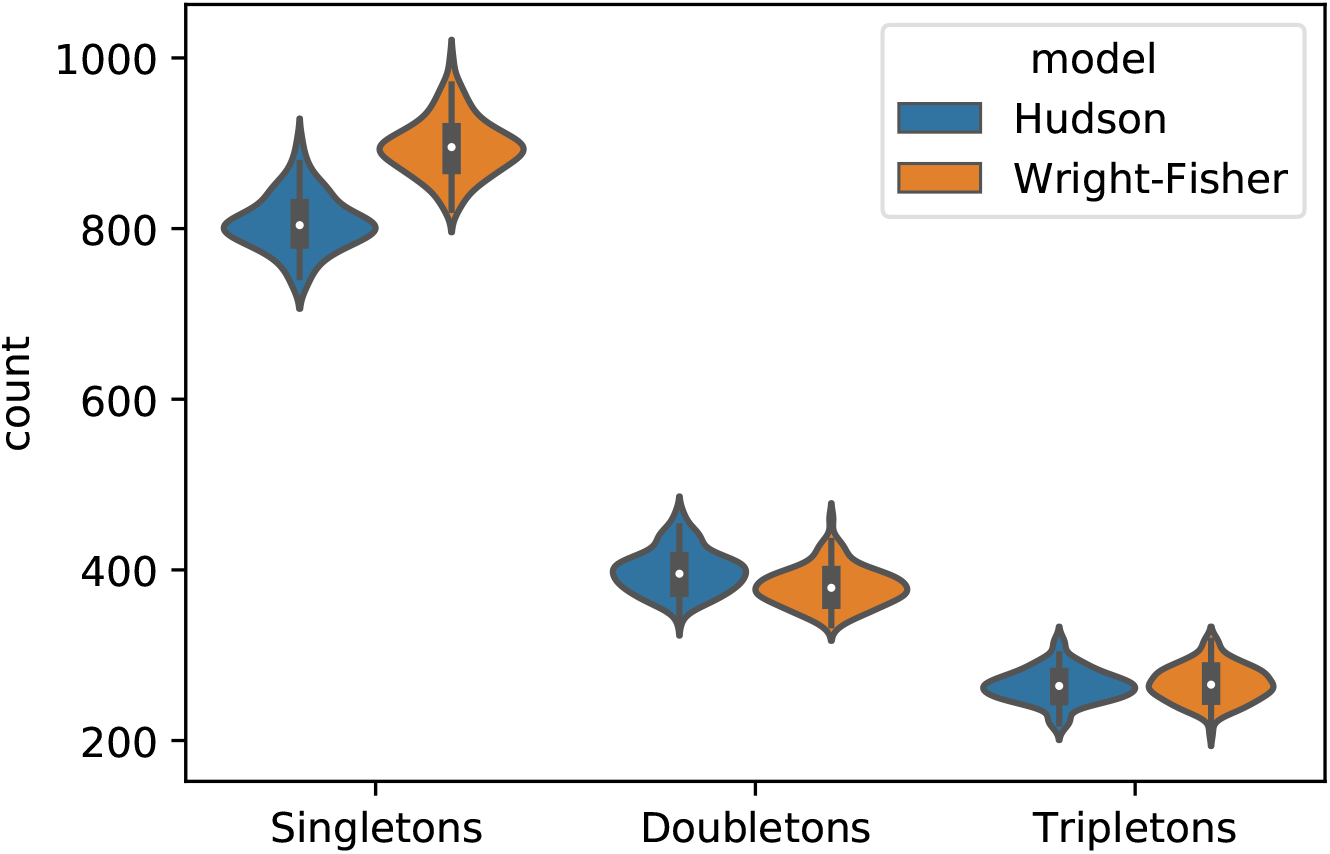
Number of singletons, doubletons, and tripletons simulated under Wright-Fisher and Hudson coalescent models. A 1Mb region was simulated 100 times in 20,000 haploid lineages in a diploid population of 10,000 individuals.

**Figure S2:**
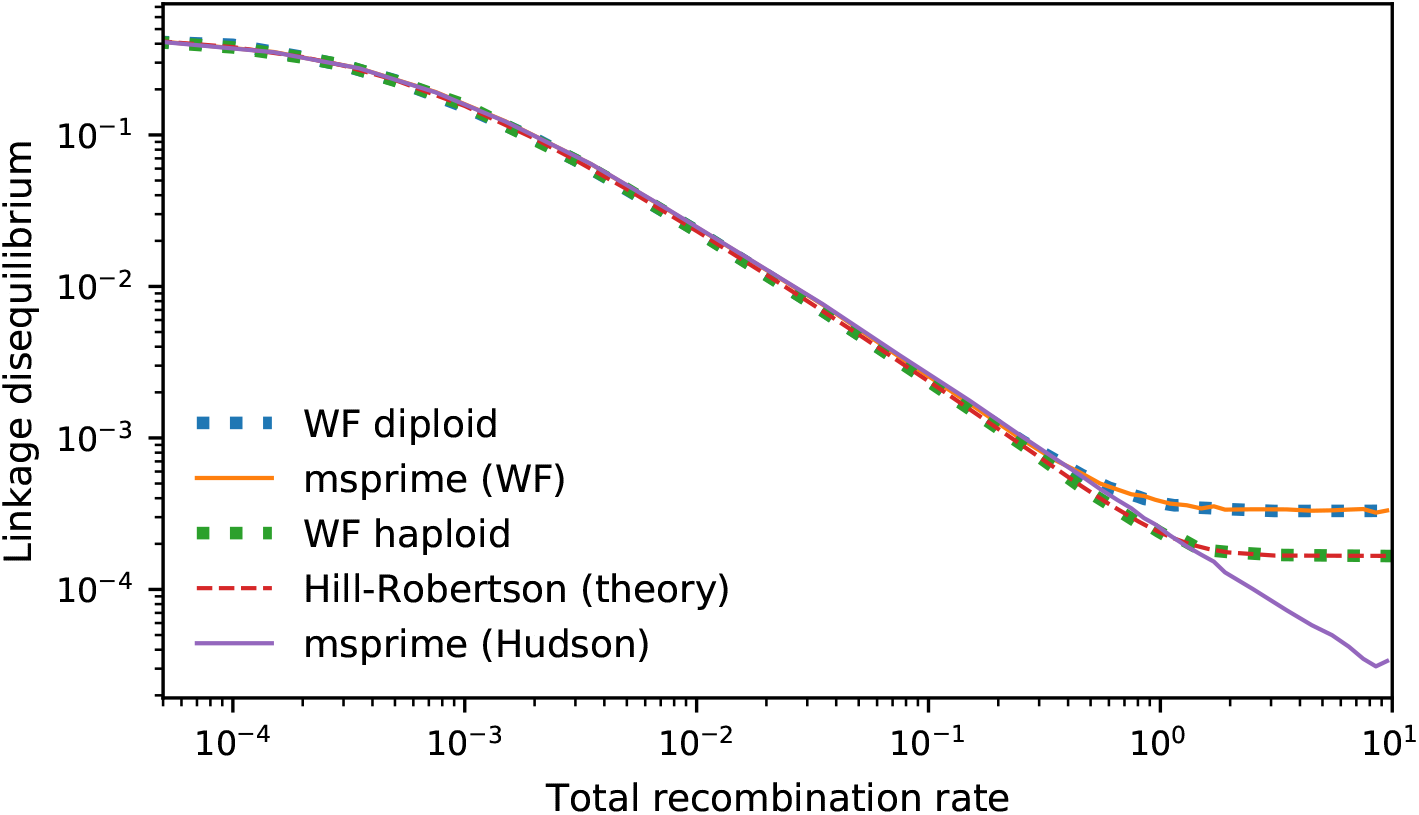
Linkage disequilibrium as measured by 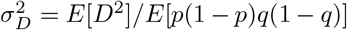 under different simulation and theory models [23]. Simulations were carried out with population size *N* = 1000 at steady state demography for a single 10M chromosome. At fully unlinked loci, the expected value of 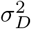 is 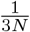 in a diploid model and 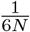 in a haploid model [24].

**Figure S3:**
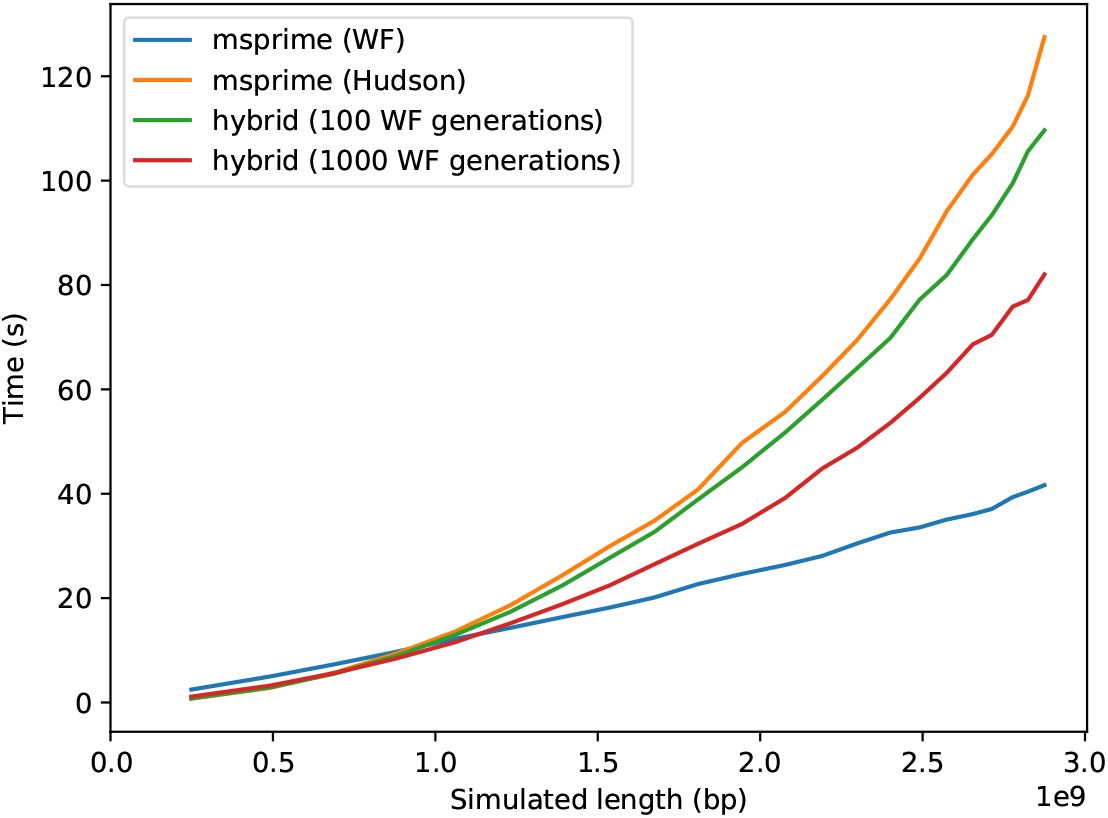
Computation time of Hudson coalescent, Wright-Fisher, and hybrid models with 100 and 1000 Wright-Fisher generations before switching to the coalescent. Simulations contain from 1 to 22 chromosomes of realistic lengths, using the method described in Section 4.1, in 500 haploid samples within a diploid population of size 500.

**Table S1:** Relative difference in mean number of singletons, doubletons, and tripletons under the Wright-Fisher (*N_W F_*) and Hudson (*N*_*H*_) models, from data shown in Figure S1. These results closely match those presented in [13].

| Frequency | $\frac{N_{WF} - N_H}{N_{WF}}$ |
| --- | --- |
| Singletons | 0.099131 |
| Doubletons | -0.047253 |
| Tripletons | 0.010092 |

